# Comparative modeling reveals the molecular determinants of aneuploidy fitness cost in a wild yeast model

**DOI:** 10.1101/2024.04.09.588778

**Authors:** Julie Rojas, James Hose, H. Auguste Dutcher, Michael Place, John F Wolters, Chris Todd Hittinger, Audrey P Gasch

## Abstract

Although implicated as deleterious in many organisms, aneuploidy can underlie rapid phenotypic evolution. However, aneuploidy will only be maintained if the benefit outweighs the cost, which remains incompletely understood. To quantify this cost and the molecular determinants behind it, we generated a panel of chromosome duplications in *Saccharomyces cerevisiae* and applied comparative modeling and molecular validation to understand aneuploidy toxicity. We show that 74-94% of the variance in aneuploid strains’ growth rates is explained by the additive cost of genes on each chromosome, measured for single-gene duplications using a genomic library, along with the deleterious contribution of snoRNAs and beneficial effects of tRNAs. Machine learning to identify properties of detrimental gene duplicates provided no support for the balance hypothesis of aneuploidy toxicity and instead identified gene length as the best predictor of toxicity. Our results present a generalized framework for the cost of aneuploidy with implications for disease biology and evolution.

## Introduction

Aneuploidy, when cells carry an abnormal number of one or more chromosomes, can produce different outcomes depending on the environmental and cellular context. On the one hand, aneuploidy is broadly considered deleterious. Amplification of most human autosomes is lethal except for trisomy of chromosome 21 that causes Down syndrome (DS)^1^. The deleterious effects of chromosome duplication can also be seen at the cellular level in most organisms^2,3^. On the other hand, aneuploidy is often beneficial during evolution. Chromosome amplifications are frequently selected in drug-resistant human pathogens and represent a major source of drug evasion^4,5^. Furthermore, aneuploidy is observed in ∼20% of non-laboratory *S. cerevisiae* isolates^6–9^ and is associated with adaptive traits in natural and industrial environments^10–15^. Aneuploidy is also found in >88% of cancers: tumors with high levels of aneuploidy display poorer patient prognosis, respond less well to treatment, and have a higher rate of relapse^16,17^. Recent studies show that specific chromosome amplifications underlie these benefits^17–21^. Thus, aneuploidy can be a fast route to adaptation in a changing environment. Whether cells can use aneuploidy to evolve to a new environment depends on the balance between aneuploidy cost and potential benefit – if the benefit under the conditions at hand outweighs the cost, aneuploidy will be maintained.

However, a major limitation in predicting the impact of aneuploidy is that we lack a mechanistic understanding of why aneuploidy is deleterious under optimal conditions, especially in the case of chromosome amplifications. Previous studies showed large mammalian chromosomes transformed into yeast and lacking coding potential do not incur the same fitness cost as duplicating native chromosomes, strongly implicating protein-coding sequences as a major contributor contributor^22–24^. Two mutually exclusive models have been proposed to explain the inherent cost of duplicating chromosomes (herein referred to as aneuploidy). On one end of the spectrum is what we refer to as the Genic Load model, in which aneuploidy cost is driven by the burden of making extra gene products, independent of their functions or properties^25,26^. Multiple studies, from yeast to mammals, suggest that larger chromosomes with more genes incur a larger cost^2,6^. In yeast, chromosome length and gene number negatively correlate with the growth defect of aneuploid strains^27,28^ and with the number of aneuploid strains found outside of the lab, which presumably reflects the strength of negative selection^6,9,29^. The magnitude of that correlation varies for different studies, which analyze incomplete sets of strains often isolated from multiple sources. In another study, the impact of large segmental duplications on yeast growth was partly correlated with the number of genes in those segments, although discrepancies were identified^30^.

On the other end of the spectrum is the Driver Gene model that predicts that aneuploidy toxicity is due to a handful of dosage-sensitive genes encoded on each chromosome. This model prevails in the study of DS, where research often focuses on one or a few specific genes on human chromosome 21^31,32^. In yeast, one of the most striking examples of a driver gene is thought to be beta-tubulin *TUB2* encoded on ChrVI (Chr6): Chr6 duplication is only viable in the presence of other chromosome duplications that encode Tub2 interacting proteins; these chromosome amplifications occur spontaneously when *TUB2* is duplicated on a plasmid^33,34^. In cancer cells, the frequency of segmental gains and losses found in the Cancer Genome Atlas database can be partially modeled by the number of tumor suppressors and oncogenes amplified in those regions^35–37^. These models are based on <8% of genes scored at the time as tumor suppressors and oncogenes, suggesting that only a subset of human gene amplifications contribute a major impact. Other recent studies provide evidence for a mixed model of aneuploidy cost. For example, Keller et al. analyzed a suite of segmental chromosome amplifications in yeast and showed that fitness cost partially correlated with the length of the amplification; however several outliers implicated that other effects must be at play^30^. A major limitation in distinguishing any of these models is a lack of systematic study measuring the cost of each chromosome’s duplication in a controlled environment and then modeling the mechanistic basis for that cost.

Both of the models above are compatible with a prominent view of deleterious effects known as the Balance Hypothesis. This hypothesis posits that duplication of genes encoding proteins with many protein interactions or that participate in multi-subunit protein complexes can produce stoichiometric imbalance, disrupting protein interaction networks and causing downstream stress on protein folding, degradation, and management known as proteostatic stress^38–40^. Proteostasis stress can be exacerbated by an increased burden produced by many gene amplifications, overloading cellular machineries^2,23,41,42^. Indeed, yeast aneuploids are sensitive to conditions that interfere with proteostatic functions including protein translation, folding, and degradation^2,23,41–43^. However, these models are heavily influenced by results from a laboratory strain of yeast, W303, that is highly sensitized to chromosome duplication. The genetic basis for this sensitivity is a hypomorphic variant of RNA-binding protein, Ssd1, that is required for yeast to tolerate extra chromosomes^44^. Most non-laboratory strains studied to date are significantly more tolerant of chromosome amplification, although a detectible fitness cost remains, raising questions about the cost and effect of aneuploidy in more representative non-laboratory strains^6,7,9^.

In this study, we used comparative modeling and molecular validation to distinguish and quantify models of aneuploidy cost in a natural oak-soil isolate of *S. cerevisiae* YPS1009, with and without *SSD1*. In doing so, we leveraged a pooled library of cloned genes to measure the cost of duplicating each gene individually. Our results indicate that a multi-factorial model incorporating the additive cost of individual gene duplications on each chromosome, plus the impact of several noncoding RNA classes, explains a large proportion of aneuploid growth defects. Surprisingly, we found no evidence for the Balance Hypothesis in aneuploidy cost and propose that yeast cells have evolved to manage mere duplication of most genes. We used machine learning approaches to identify other features associated with deleterious single-gene duplication. Surprisingly, the most impactful feature predicting the fitness effect of a gene’s duplication is its length, since deleterious genes are on average significantly longer than non-deleterious genes. Together, our results raise important considerations regarding the effects of gene and chromosome amplification.

## Results

### Fitness costs of chromosome duplication vary by chromosome

We began by generating a panel of haploid strains in the oak-soil YPS1009 background in which each chromosome is duplicated. We used the method of Hill et al.^45^ to generate aneuploid cells by integrating a galactose-inducible promoter facing each centromere. Cells were shifted to galactose medium for one generation to induce transcription, which blocks kinetochore assembly and thus causes chromosome retention in the mother cell during mitotic division (see Methods). We generated aneuploid strains in which each of 15 out of the 16 yeast chromosomes is duplicated. The exception was Chr6, proposed previously to be lethal due to amplification of tubulin *TUB2*^33,34,46,47^. Most of the chromosome duplications were stable over many generations (see Methods). We generated a comparable strain background that was sensitized to aneuploidy through the deletion of *SSD1*^44^. We were unable to isolate a *ssd1Δ* strain with Chr16 duplicated, suggesting that this specific chromosome duplication is not viable in this strain background without Ssd1.

The strain panel affords an opportunity to sensitively measure the fitness cost of aneuploidy under standardized conditions. We measured the growth rate of wild-type and *ssd1Δ* strains in the panel, in biological quadruplicate. Not surprisingly, different chromosome duplications inflict different levels of fitness defects (Fig. 1A). We observed a range of growth rates, from 96% of the euploid growth rate for duplication of Chr3 (the 2^nd^ smallest chromosome) to 65% for Chr15 duplication, which falls among the larger chromosomes but is not the largest in size or gene content. These results already highlight an imperfect relationship between chromosome size and its fitness cost. Ssd1 was previously shown to be important for some chromosome duplications in multiple wild strains^44^, but the breadth of its impact on other chromosomal aneuploidies was not previously known. We discovered that 9 of the 15 aneuploids (60%) incurred significantly greater growth defects in the *ssd1Δ* background (Fig. 1A). Most of the other chromosomes were also more deleterious in the *ssd1Δ* strain but missed the threshold for statistical significance. Thus, Ssd1 is important for tolerating most chromosome duplications, with greater impacts for chromosomes that cause a greater defect in wild-type cells.

**Figure 1.**
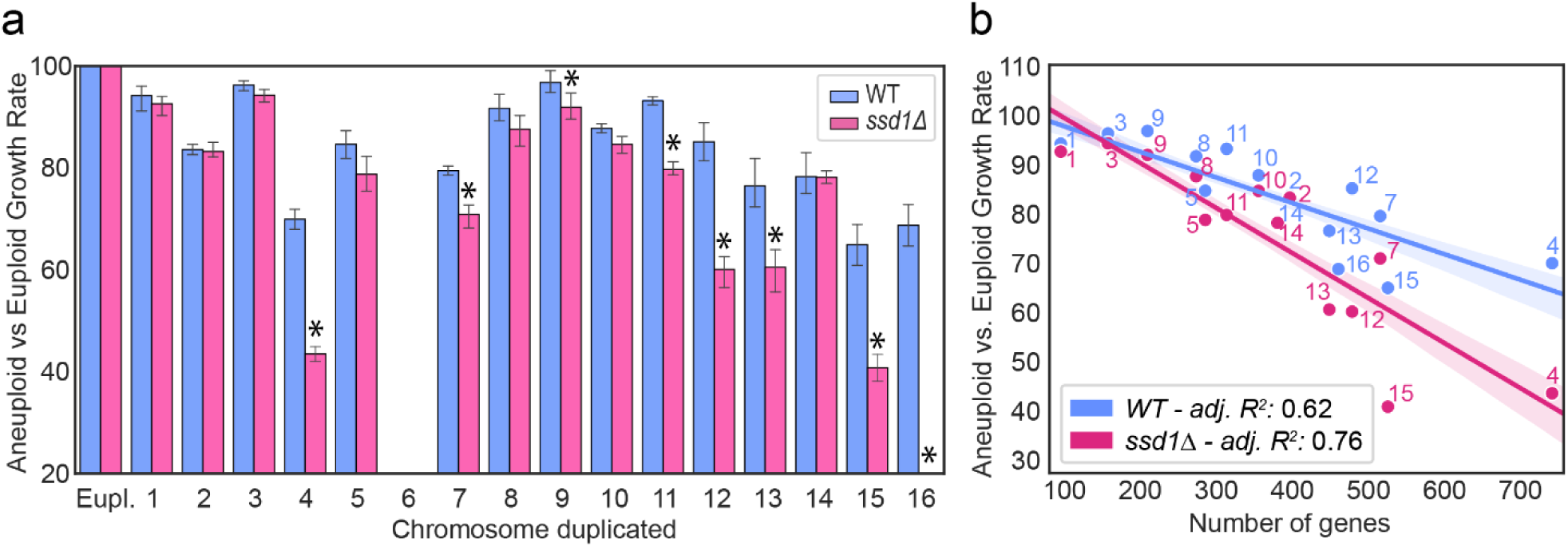
Chromosome duplications inflict variable fitness costs in wild-type and *ssd1Δ* cells. (A) Average and standard deviation (n=4) of aneuploid growth rates relative to isogenic euploid. All *SSD1+* (‘WT’, blue) aneuploids grew slower than the euploid (p<0.05, replicate-paired T-test); *ssd1Δ* aneuploids that grew significantly slower than their wild-type aneuploid equivalent are indicated with an asterisk (p<0.05, T-test). (B) Mean relative growth rate of each aneuploid strain (numbered by duplicated chromosome) relative to the isogenic euploid plotted against the number of genes per amplified chromosome. Ordinary least squares regression with 95% confidence interval shaded and adjusted R^2^ indicated in the box.

### Genic load partly explains the fitness costs of chromosome duplication

With the fitness costs of each chromosome duplication in hand, we developed mathematical models to understand the determinants of aneuploidy toxicity. For optimal modeling, we first sequenced the YPS1009 genome using long-read sequencing combined with short-read polishing. This produced a high-quality genome of 7,362 annotated genes and non-genic features across 16 assembled chromosomes (see Methods).

We began by calculating the linear relationship between the relative fitness cost measured for each chromosome duplication (taken as the aneuploid versus euploid growth rates) and the number of genes per chromosome (Model 1), which in yeast is highly correlated with the chromosome length (R^2^ = 0.99). Excluding Chr6 that could not be generated, the fit for the remaining chromosomes explains 62% (adjusted R^2^ = 0.62) of the variance in relative fitness costs. Thus, the number of genes per chromosome alone explains a significant proportion of the variance of aneuploids fitness cost (Fig. 1B), confirming previous implications in various organisms^6,9,27,28^. The fit was even higher for *ssd1Δ* strains, explaining 76% of the variance in fitness costs of cultivatable chromosome duplications (Fig. 1B). The increased slope reflects the stronger fitness costs in *ssd1Δ* aneuploids, suggesting that, in the absence of Ssd1, cells are more sensitive to the genic load.

### The additive effect of single gene duplications accurately models whole chromosome gain

An open question in the aneuploidy field is the degree to which specific genes on each duplicated chromosome contribute to the fitness cost of aneuploidy. We therefore set out to determine the fitness impact of duplicating each gene individually in the YPS1009 euploid strain using a single-copy gene duplication library. Each centromeric plasmid contains one yeast gene with its native upstream and downstream regulatory sequence along with a unique DNA barcode for tracking^48^. To measure the fitness cost of duplicating each gene individually, we transformed euploid YPS1009 with the pooled library and grew competitively for ten generations, taking the log_2_(fold change) in barcode abundance after competitive outgrowth as the relative fitness cost (see Methods). Genes whose barcode abundance significantly decreased during growth are considered detrimental to fitness, while those whose frequency increased are considered beneficial. Out of the 4,369 YPS1009 yeast genes for which fitness could be measured, 25.5% were beneficial and 28% were detrimental (FDR < 0.05, Fig. 2A, Fig. S1). Because genes with noisy measurements are statistically insignificant but can have artificially skewed mean measurements, we replaced insignificant scores (FDR > 0.05) with the mean cost of all measured genes (log_2_ value of -0.33, see Methods). Hence, genes without a significant effect are considered to have a mild negative impact. We then computed the fitness cost of each chromosomal duplication (Chr. cost) as the additive fitness cost of all genes encoded on that chromosome (*i.e.* the sum of log_2_ fitness effects).

**Figure 2.**
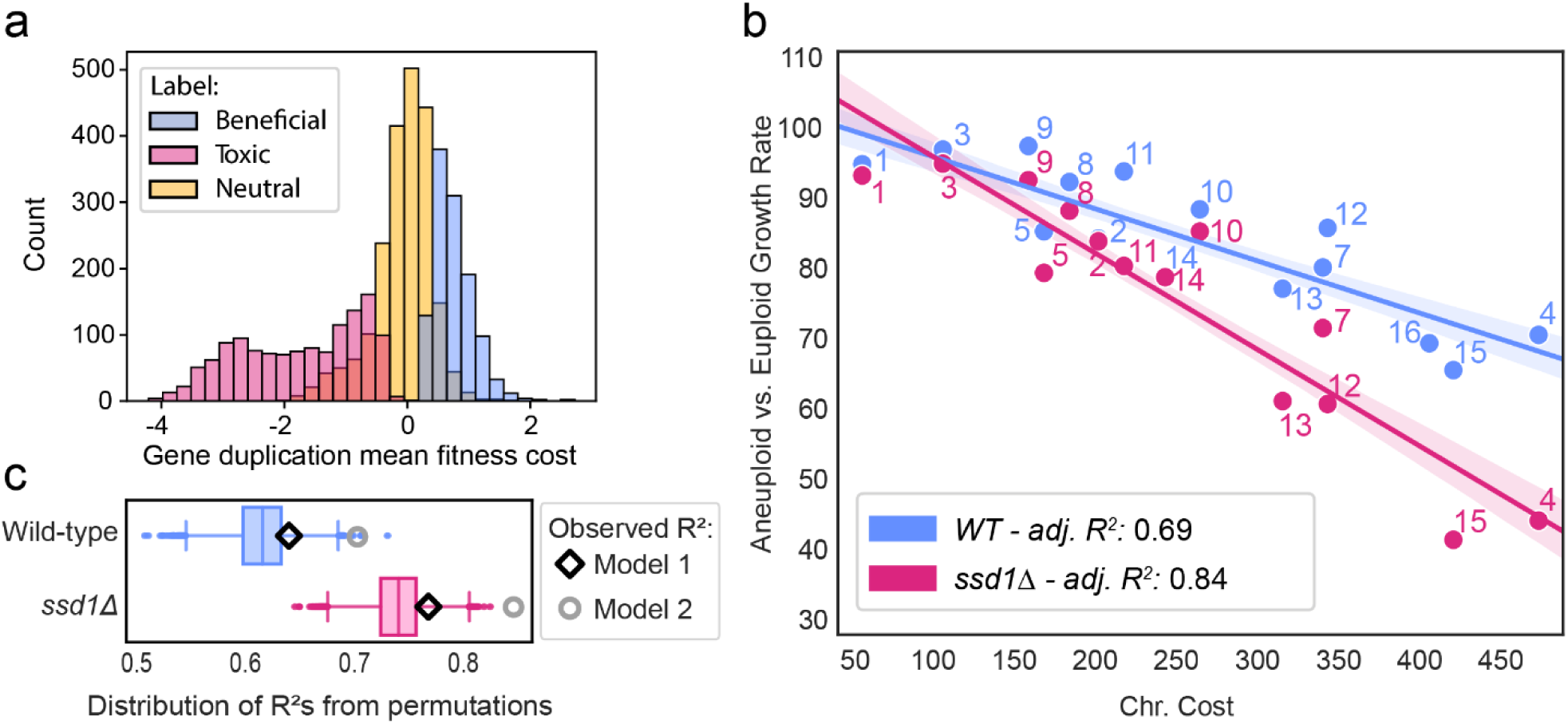
Considering gene-specific fitness costs improves the modeling. (A) Distribution of log_2_ fitness scores for single-gene duplications for gene groups in the key. (B) Linear fit of the mean relative growth rate as in Fig 1 plotted against the sum of the log_2_ fitness costs for genes encoded on each chromosome (‘Chr. Cost’). (C) Distribution of R^2^ values from 10,000 random permutations of gene fitness scores affiliated with each chromosome. The observed adjusted-R2 values for Model 1 and Model 2 are shown for each strain panel.

The additive model of single-gene costs (Model 2) significantly improved the fit compared to Model1 that considers only the number of genes per chromosome (Adjusted R^2^ for wild-type = 0.69, *ssd1Δ* = 0.84, Fig. 2B). The improvement in the fit was highly significant as assessed in two ways. First, a nested model that included both the number of genes per chromosome and the additive-gene cost (normalized to chromosome gene number) improves the fit, since both factors are significant (p<3.9×10^-2^, likelihood-ratio test, see Methods). Second, the observed fit for Model 2 was better than nearly all of the 10,000 random permutations of gene fitness costs (preserving the number of genes per chromosome in each trial). Out of 10,000 permutations, only 4 met the observed Model2 fit for wild-type aneuploids (p = 0.0004) and none for the *ssd1Δ* strains (p < 0.0001, Fig. 2C). Together, these results show that the identity of duplicated genes has an important contribution to the cost of aneuploidy and is predictive of the fitness effect of whole chromosome duplication. Interestingly, the observed fit for Model 1 that simply counts the number of genes per chromosome was better than 88% and 82% of random trials for the wild type and *ssd1Δ* strains, respectively, which were close to statistical significance (p = 0.17 for wild-type, p = 0.11 for *ssd1Δ*). This raises the intriguing possibility that fungal evolution has optimized gene content on each chromosome to minimize the cost of chromosome duplication, which is relatively frequent in yeast. Regardless, these results show that the combination of single-gene fitness effects is predictive of the fitness effect of whole chromosome duplication (see Discussion).

We devised an independent experimental approach to test the models using strains carrying two chromosome duplications. These dual-chromosome duplications were not stable in *ssd1Δ* cells, and thus we focused on *SSD1+* strains. Those with multiple chromosome duplications grew slower than corresponding single-chromosome duplication strains, as expected (Supplemental Fig. S2A). We assessed the variance in growth rates of dual-chromosome duplications explained by the models trained on single-chromosome duplications. Indeed, Model 2 was significantly better (adj. R^2^ = 0.54) than Model 1 (adj. R^2^ = 0.34, Fig. S2B-C). Thus, the model does not overfit the training data and instead shows that the cost of chromosome duplication is significantly influenced by the suite of genes encoded on each chromosome.

### Beneficial gene duplications alleviate the cost of chromosome duplication

Studies in cancer cells suggested that beneficial oncogenes on amplified chromosomes counteract tumor suppressors on the same segments^35,36^. We wondered if genes whose duplication is beneficial to YPS1009 are important for aneuploidy fitness. To test this, we excluded beneficial genes from the Chr. cost, which decreased the model performance (adjusted R^2^ of 0.67 for wild-type and 0.80 for *ssd1Δ* compared to 0.69 and 0.83, respectively, for Model 2). The contribution of beneficial genes is statistically significant in a nested model in which their additive effect was added as a separate feature (p-value = 4.7 x10^-2^, likelihood test). Hence, genes that are beneficial when duplicated in isolation contribute to aneuploidy fitness, likely because they collectively counter some of the aneuploidy fitness cost.

### Non-coding features contribute to aneuploidy fitness effects

While it is clear that gene fitness costs explain much of the cost of chromosome duplication, non-coding features could also contribute. We therefore compiled a set of non-genic features per chromosome based on the YPS1009 genome sequence and used Lasso regression to identify additional features that improve predictions. The input set included the number of small nucleolar (sno)RNAs, tRNAs, other non-coding (nc)RNAs, autonomous replicating sequences (ARS), retrotransposons, and long-terminal repeats (LTR), all normalized by the total number of features encoded on each chromosome (see Methods). Aside from LTR and retrotransposon numbers, most of the features were not confounded by co-variation (Supplemental Fig. S3).

We used a bootstrap-Lasso (Bolasso-S, Bach 2008) approach to select features that contribute significant explanatory power to the modeling of measured aneuploidy growth defects, selecting from non-genic features as well as Model 2 genic costs per chromosome. Features selected by Lasso regression in 90% of 1000 Bootstrap trials (Lasso alpha factor = 0.7, see Methods) were retained and incorporated into multi-factorial Model 3. For both wild-type and *ssd1Δ* models, Lasso chose the Chr. costs from Model 2 as the most impactful factor but also the normalized number of snoRNAs per chromosome as deleterious to fitness and the normalized number of tRNAs per chromosome as beneficial (Fig. 3A). In addition to these features, the method also chose the normalized number of retrotransposons as a deleterious predictor only for the *ssd1Δ* strain. All selected features were significant (Chi-Square test, Fig. 3A). Remarkably, the multi-factorial Model 3 explains 74% of the growth rate variance for wild-type and 94% for *ssd1Δ* aneuploids (Fig. 3B). When the trained models were assessed on dual-chromosome duplication strains, Model 3 improved the predictions compared to Model 2 (adjusted R^2^ = 0.7 compared to 0.54 for Model 2, Supplemental Fig. S2D).

**Figure 3.**
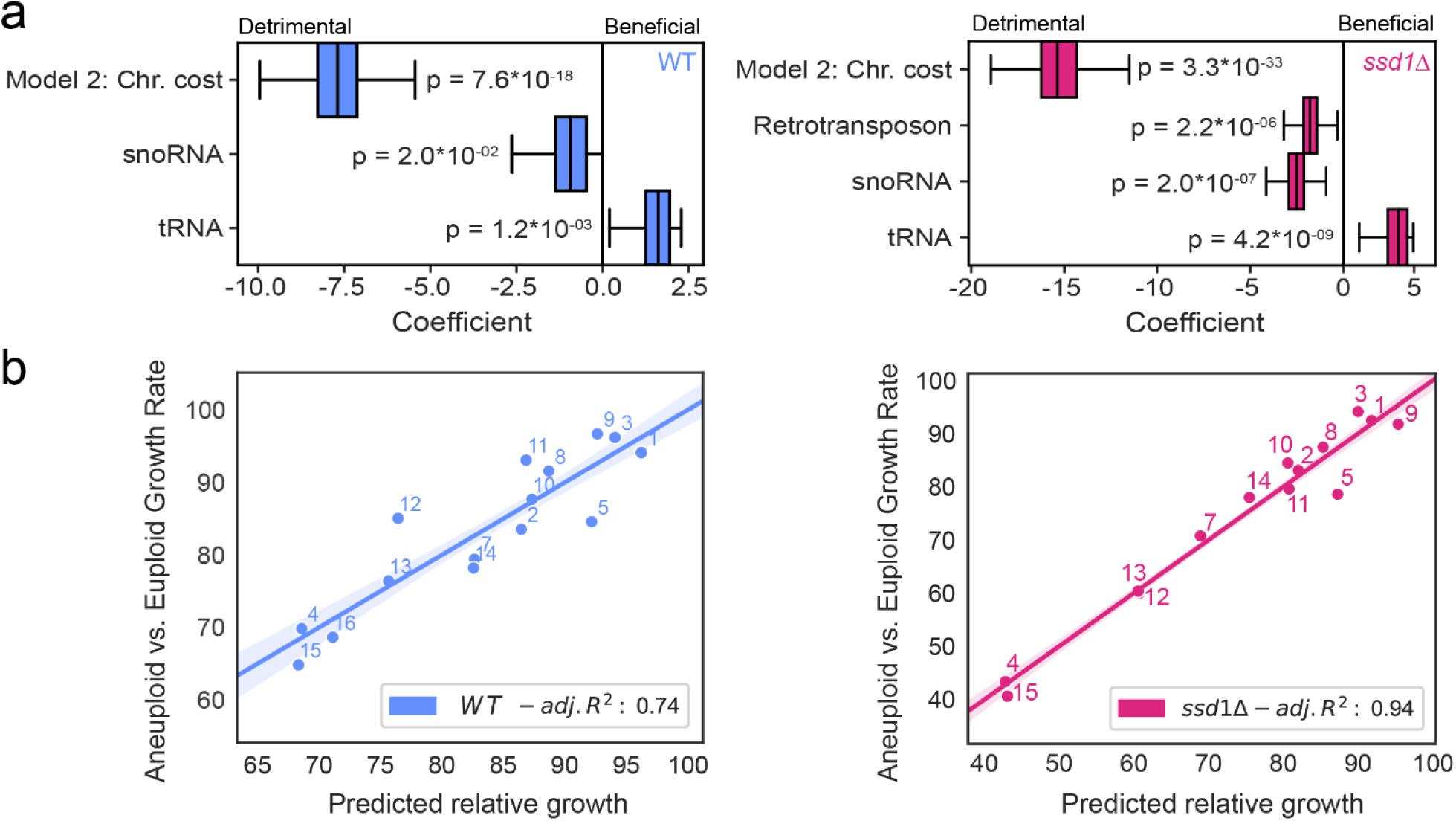
A multi-factorial model best explains the costs of chromosome duplication. (A) Distribution of coefficients obtained from 1000 Lasso regression bootstrap iterations. Only features exhibiting non-zero weights in more than 90% of bootstrap resamples are depicted. The Likelihood-ratio test’s p-values for each selected feature for the wild-type (blue) and *ssd1Δ* (pink) regression models are displayed on the figures. (B) Linear fit of the mean relative growth rates as in Fig 1 against Model 3 predictions (using significant features for each strain as shown in A). The adjusted R-squared value is indicated in the lower right corner.

### Imbalanced duplication of snoRNAs is detrimental

The Lasso predictions above improve the modeling, but is the model correct? We set out to experimentally verify several of the model predictions. We first tested the predicted deleterious impact of duplicating snoRNAs. snoRNAs guide catalytic modifications of other RNAs, such as ribosomal rRNAs and tRNAs. snoRNAs can be split into C/D box snoRNAs that direct 2’-hydrolyl methylation of their RNA targets, and H/ACA box snoRNAs involved in pseudouridylation^50^. The two groups were combined into one for modeling given their relatively small numbers in the genome (45 C/D and 29 H/ACA). We cloned 7 C/D snoRNAs present in an array on Chr13 or 7 H/ACA snoRNAs from a single region on Chr15 onto centromeric plasmids (see Methods). Duplication of both snoRNA cassettes significantly reduced growth of the euploid strain, validating that duplication of these cassettes is indeed deleterious (Fig. 4A). Furthermore, the growth rates of haploid YPS1009 carrying duplications of Chr4 or 15 were also reduced upon duplication of these snoRNAs (despite missing the significance threshold in one case, Fig. 4A).

**Figure 4.**
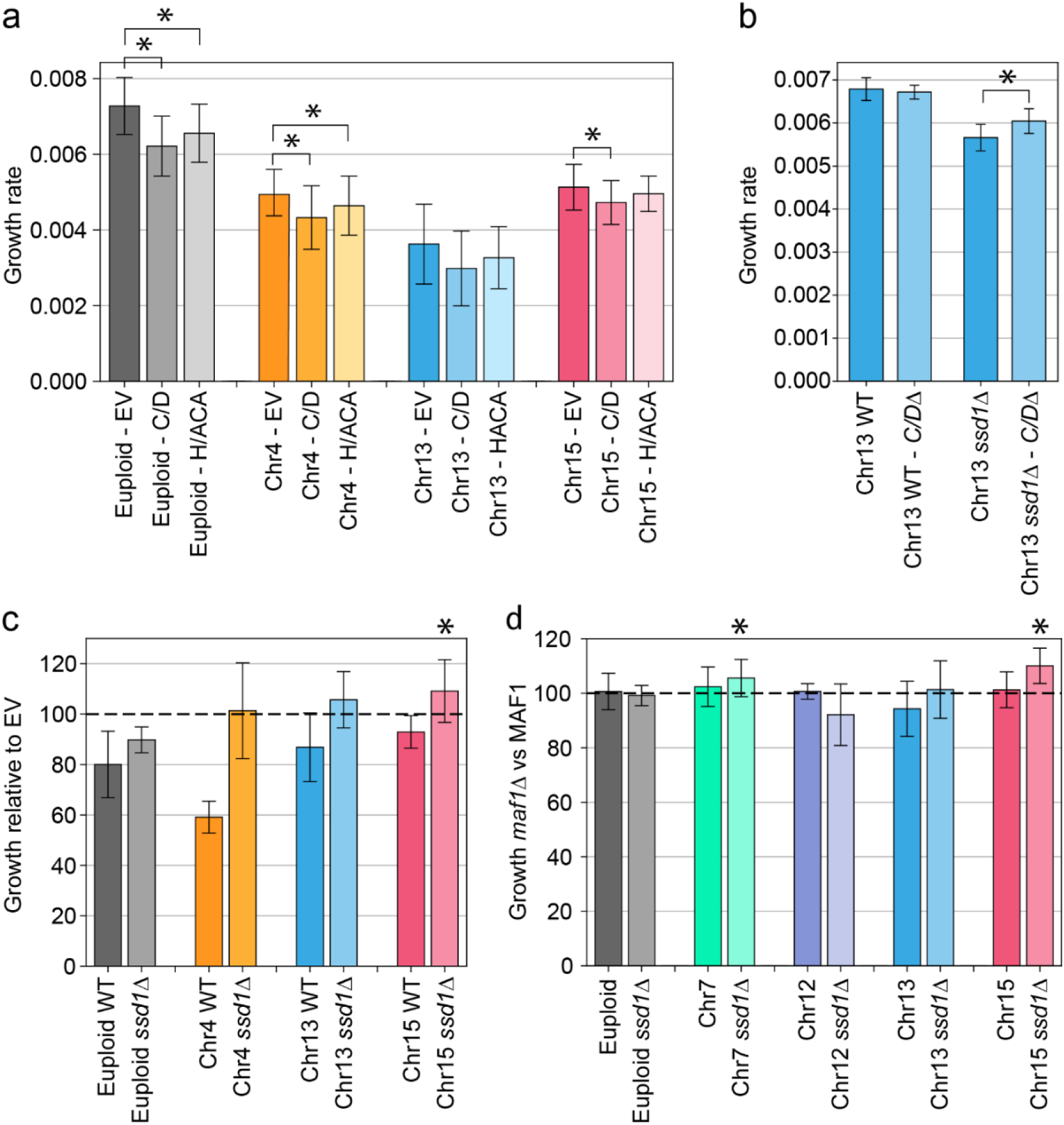
Duplication of select snoRNAs and tRNAs contributes to aneuploidy fitness. (A) Average and standard deviation of growth rates of strains containing the empty vector (EV) or plasmids encoding either 7 C/D box snoRNAs or 7 H/ACA snoRNAs as described in the text (*, p <0.05, replicate-paired T-test versus empty vector). (B) Average and standard deviation of growth rates of Chr13 aneuploids with or without restoring 7 C/D box snoRNAs copy number to euploid levels. (*, p <0.05, replicate-paired T-tests). (C) Average and standard deviation of relative growth rates of strains harboring Chr 12-tRNA cassette versus strain with the empty vector (*, p < 0.01, replicate paired T-tests, between each aneuploid and the corresponding euploid). (D) Average and standard deviation of relative growth rates of each strain in the *maf1Δ* versus *MAF1+* background (*, p < 0.05, replicate-paired T-tests between *MAF1* and *maf1 Δ*).

Reciprocally, if snoRNAs contribute to aneuploidy toxicity, then restoring to euploid copy number should partially alleviate the aneuploidy fitness costs. In that aim, we deleted from one of the Chr13 copies a segment of 6 out of its 9 C/D snoRNAs (see Methods). Although there was no significant effect in the wild type, deleting the extra C/D snoRNA copies from the *ssd1Δ* Chr13 aneuploid strain significantly improved its growth rate. The increased sensitivity of *ssd1Δ* aneuploids may provide more power to detect improvements than in the wild type, where snoRNA imbalance was also predicted to be deleterious. Nonetheless, together these results confirm that snoRNA duplication is deleterious to both euploid and aneuploid cells and contributes to the cost of chromosome duplication in at least the *ssd1Δ* background (see Discussion).

### Increasing tRNA copy number benefits *ssd1Δ* aneuploid cells

Model 3 above predicts that chromosomes with more tRNAs are less deleterious than otherwise predicted. We tested this in several ways. First, we introduced a plasmid carrying 21 tRNAs encoded on Chr12^51^ into the YPS1009 euploid and a subset of aneuploid strains. The tRNA plasmid decreased proliferation in the euploid and Chr4 aneuploid wild-type cells, indicating that an imbalanced set of these tRNAs is deleterious (Fig. 4C). However, their duplication had a less detrimental effect in the other aneuploids, especially strains lacking *SSD1*. In fact, duplication of these tRNAs was beneficial to varying degrees in *ssd1Δ* aneuploids with Chr13 and Chr15 duplications.

As an alternative approach, we assessed the effect of upregulating all tRNAs by deleting the RNA polymerase III repressor, Maf1. *MAF1* deletion leads to an accumulation of tRNAs^52^, which we confirmed (Supplemental Fig. S4). We found that *MAF1* deletion improved growth rates for Chr7 and Chr15 aneuploids in the *ssd1Δ* background (p-value < 0.05, Fig 4D). Although the effects were somewhat mixed, these results suggest that several aneuploidy-sensitized *ssd1Δ* strains benefited from extra tRNAs but that the effect could be specific to certain chromosomes or tRNAs (see Discussion).

### Machine learning implicates properties common to duplication-sensitive genes

Although the additive cost of gene duplication explains a significant proportion of the cost of aneuploidy, some gene duplicates are more deleterious than others. To further explore this, we sought properties that are predictive of deleterious genes. We focused on 1,177 genes scored as deleterious when duplicated in euploid YPS1009 (FDR < 0.05), compared to 3,028 genes whose duplication was neutral or beneficial (FDR > 0.05, herein referred to as ‘neutral’). Consistent with other studies using gene duplication libraries^53–57^, we found only a handful of functional terms enriched in the deleterious group, including several categories linked to cell-cycle regulation. We next compiled a list of 120 gene and protein properties and selected those that differentiated the deleterious gene duplications from the neutral set (Wilcoxon rank sum test, Supplemental Fig. S5A, Table S1-S2). The group of deleterious genes displayed a slightly higher proportion of intrinsically disordered regions, marginally more phosphorylated sites, a higher proportion of serine residues, lower translation rates as indicated by ribosome profiling^58^, and longer length (Supplemental Fig. S6); however, several of these features are correlated with one another (Supplemental Fig. S5A), confounding interpretation. Notably, the group of genes that are deleterious when duplicated was not enriched for those encoding proteins involved in complexes or with a high number of protein-protein interactions (see Discussion).

We used a machine-learning approach to identify the most impactful gene properties and determine if their combination can accurately differentiate deleterious gene duplications from those that are neutral or beneficial (see Methods). A logistic regression classifier was trained on significant biophysical and functional enrichment (Fig. S5A, see Methods). Five-fold cross-validation revealed that the model performed relatively poorly, with a mean area under the curve (AUC) of 0.62 (Fig. 5A). Restricting the classification to the 613 most deleterious genes (bottom 15% quantile) and the 1,472 genes most confidently called neutral/beneficial (upper 65% quantile, see Methods) improved performance with an AUC of 0.713, correctly predicting 57% of deleterious gene duplications (Fig. 5A-B). Surprisingly, by far the most impactful feature in explaining deleterious genes was gene length; deleterious genes are significantly longer than neutral genes (Fig. 5C-D). A model considering only gene length had nearly equal predictive power as the more complex model (Fig. 5A). In an attempt to identify other gene properties that could in combination supplant gene length in the model, we trained a classifier without considering gene length; but the classifier performed worse (mean AUC = 0.66) than when fitted on gene length alone, and the most impactful features selected (ratio of buried residues and the presence of disordered regions) both correlate with gene length (Supplemental Fig. S5A). Thus, gene length does a better job of distinguishing the deleterious gene set than any other combination of considered features.

**Figure 5.**
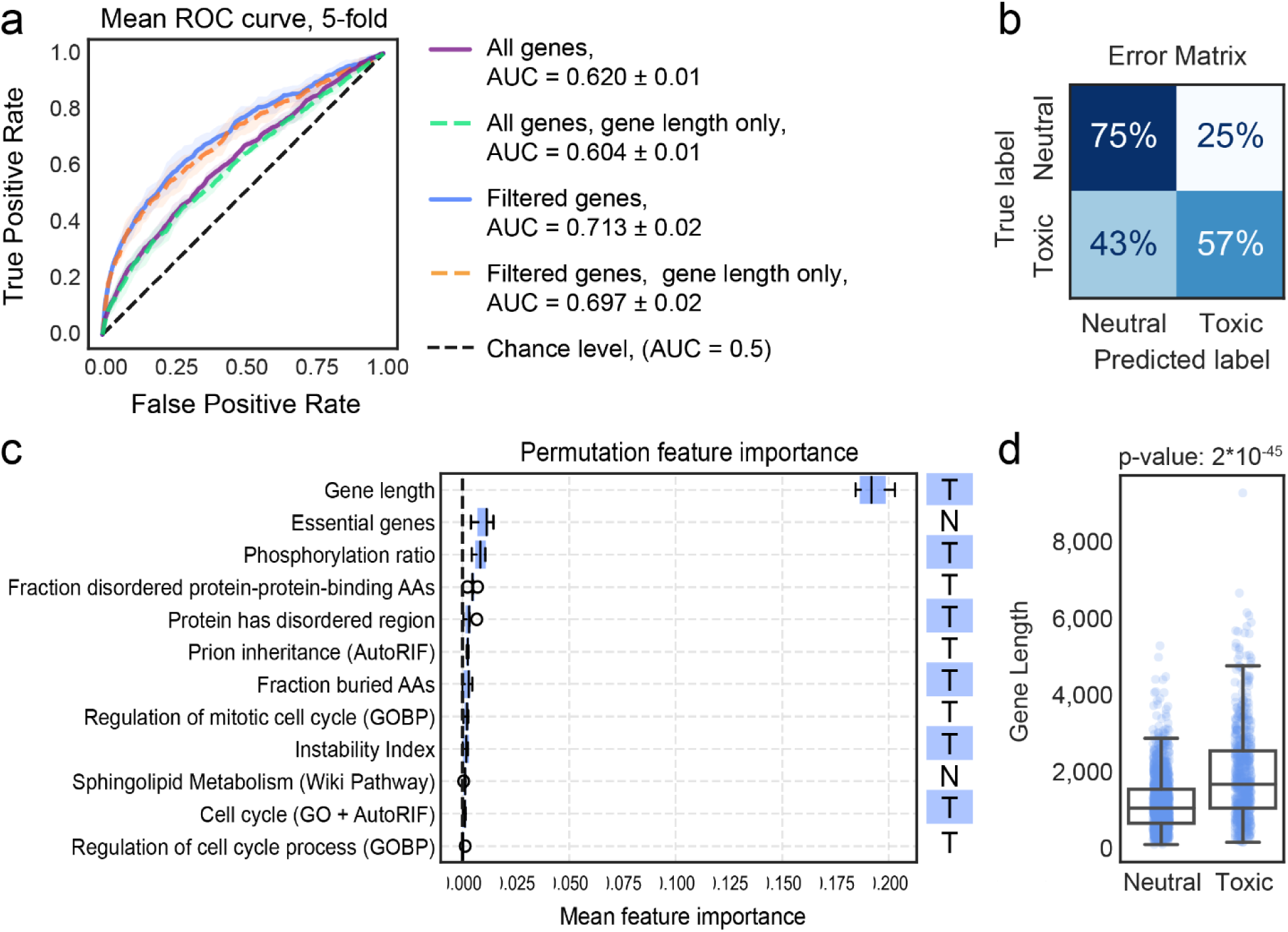
Gene length is the main predictor of deleterious gene duplications. (A) Mean ROC-curve for 5-fold cross-validation of the Logistic regression model using the top 12 features (see Methods), applied to 1,177 deleterious and 3,028 neutral gene duplications (All genes) or the restricted set of 613 substantially deleterious genes and 1,472 clearly-neutral genes (Filtered genes). Dashed, colored lines show the fit when only gene length is considered in the model. The mean Area Under the Curve (AUC) is shown in the key. (B) Error matrix shows the percent recovery of true labels by the predicted labels of the combined 5-fold cross-validation test sets. (C) Mean and standard deviation of the feature importance measured with respect to ROC-AUC gain (see Methods). Features associated with or higher in the deleterious gene duplication group are labeled with a ‘T’ while enrichment in the neutral group is indicated with a ‘N’. (D) Distribution of gene lengths for the 613 deleterious (“toxic”) and 1,472 neutral gene duplicates, p-value, Wilcoxon rank sum test.

These results were especially surprising because past work from our lab using a higher-copy library identified shared features among genes that are deleterious when over-expressed, including genes encoding proteins with many protein interactions, higher expression, intrinsic disorder, and other features^59^. We therefore applied our modeling approach to discriminate 400 genes whose higher-copy expression on a two-micron plasmid is deleterious to many strain backgrounds from genes that are neutral or beneficial in most strains (1,657 genes)^59^. This classifier was highly accurate with an AUC of 0.92, correctly predicting 82% of deleterious genes (Fig. 6A-B). Thus, the poor performance in predicting duplication-sensitive genes is not due to our methods. In fact, the model trained on the higher-copy 2-micron library performed relatively poorly when applied to the gene-duplication datasets (Fig. 6A), with an AUC of 0.68 that was once again no better than considering gene length alone. The only common predictor between the models trained on duplicated genes versus the 2-micron overexpression experiment^59^ was gene length, along with different measures of intrinsically disordered regions suggesting it as a common factor (Fig. 6C). However, the latter features had only a marginal contribution to explaining deleterious gene duplicates while being a very prominent feature for gene overexpression.

**Figure 6.**
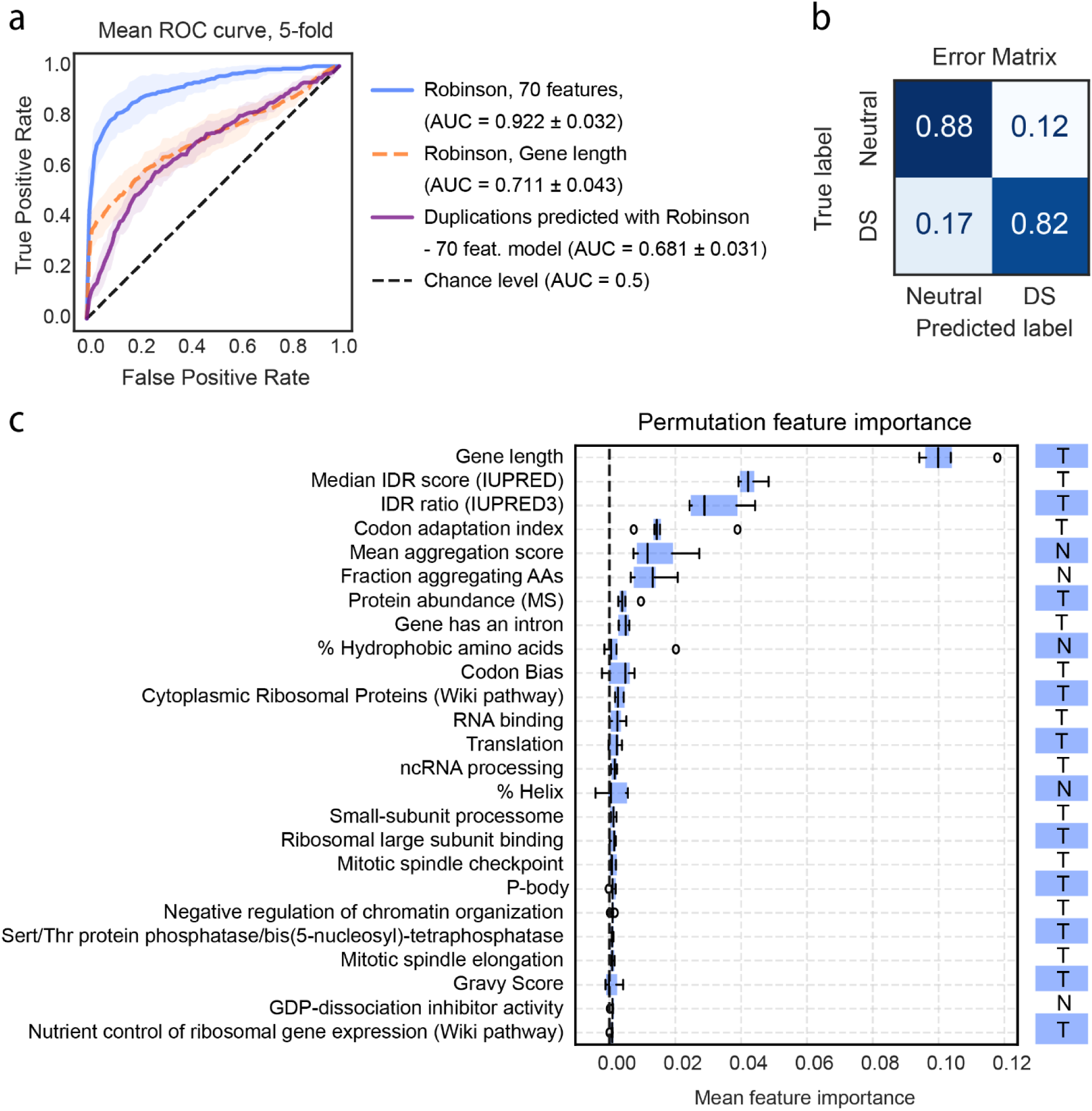
Model predictions applied to Robinson *et al.* 2-micron over-expression dataset. (A) As shown in Figure 5 but using the top 70 identified features applied to 400 commonly deleterious genes versus 1,657 commonly neutral genes based on Robinson *et al.* data^59^ (blue curve). Robinson data fit only with gene length (dashed line), or gene-duplication data from this study (“Duplications”, purple curve) fitted with the model trained on Robinson data. (B) Error matrix for Robinson *et al.* model as described in Figure 5. (C) Mean and standard deviation of the feature importance measured with respect to ROC-AUC gain (see Methods) for Robinson’s model with the top 25 features, as shown in Figure 5. A complete report of the permutation feature importance for all 70 features of the model is available in Fig.6C supplemental table.

We conclude that most biological features that account for deleterious effects when genes are over-expressed to higher levels may not be relevant for mere gene duplications. In both models, but especially in the case of gene duplication, gene length is the single best predictor of whether a gene duplication will be deleterious to strain fitness. We discuss the interpretation and implications of this result below.

## Discussion

Through systematic experimental and mathematical analysis, our results present a clarified view of the cost of chromosome duplication and the molecular properties behind it. Under standard growth conditions, the cost of aneuploidy cannot be fully explained by generic gene load nor by a handful of duplication-sensitive genes. Instead, our results quantitatively confirm previous suppositions^30,60^ that both the generalized burden of aneuploidy load coupled with combinatorial effects of the specific suite of genes and non-genic features on each chromosome explain 74-94% of the aneuploidy costs measured here. Some duplicated genes are more deleterious than others, while beneficial genes help to counteract the burden of deleterious genes on the same chromosomes. Thus, the cost of chromosome duplication is an emergent property of the affected genes and the collective burden of amplifying coding and non-coding genetic elements. Although not investigated here, it is likely that genetic interactions among genes duplicated together also contribute, albeit to a lesser extent than additive effects, perhaps explaining a portion of the 6-26% of variance not explained by our models.

Although the cost of chromosome duplication is explained by these combined effects, it is important to highlight that duplication of single genes on a chromosome can have a disproportional impact on specific phenotypes. This may explain arsenic resistance contributed by amplification of *S. cerevisiae* Chr16, which encodes arsenic resistance genes^9^, or fluconazole evasion by amplification of *C. albicans* Chr5, which encodes drug pumps and their regulators^4^. A similar implication was made for trisomy 21, by correlating specific DS phenotypes to genes amplified in subsets of people with partial-chromosomal trisomies^61,62^. These single-gene effects almost certainly contribute to chromosome-specific impacts observed for different karyotypes^3,6^. In terms of evolution, if the benefit provided by the resulting phenotypes outweighs the underlying cost of chromosome amplification, aneuploidy will be maintained. Notably, this cost-benefit analysis is heavily dependent on the environmental context, and that balance can shift with changing environments.

### The contribution of snoRNAs and tRNAs points to aneuploidy impacts on translation

Our work implicates the contribution of non-coding RNAs to the cost of chromosome duplication. The modeling predicted and experimental analysis confirmed that imbalanced expression of tested snoRNAs incurs a fitness cost in euploids and select aneuploids, whereas restoration of their balance can alleviate toxicity in the *ssd1Δ* Chr13 aneuploid. The altered abundance of specific snoRNAs can produce cellular phenotypes. For instance, overexpression of snoRNA *SNR51* in budding yeast increases binding of its target RNAs^63^. Several snoRNA-mediated modifications are found in only a portion of each substrate, such that modifications can contribute to cellular heterogeneity including ribosome functions^64^. Thus, aneuploidy-induced imbalance could change the landscape of rRNA and tRNA modifications leading to broader effects on translation^65^.

In contrast, chromosomes with more tRNAs were less toxic than the model otherwise predicted, in both wild-type and *ssd1Δ* strains, pointing to a role for tRNAs in alleviating the cost of aneuploidy. We confirmed this prediction experimentally in several sensitized *ssd1Δ* aneuploids, albeit with mixed results in the wild-type strain. We considered all tRNAs together as our relatively small dataset does not have the statistical power to test individual tRNA contribution, but different tRNA duplications may differentially benefit different chromosome amplifications. What could be the reason? The abundance of specific tRNAs correlates with the frequency of their cognate codons in the transcriptome, since higher abundance of those tRNAs facilitates translational efficiency through those codons. In fact, tRNA pools can shift composition to accommodate a changing transcriptome^66^. In recent years, tRNAs overexpression emerged as an important feature of cancer^67–69^, since upregulation of specific tRNAs increases translation of transcripts enriched for their cognate codons, thereby promoting metastasis^70,71^. Thus, the benefit of specific chromosome arm gains could be partially linked to specific tRNA duplications.

The implication of snoRNAs and tRNAs adds to a growing body of evidence that aneuploids may have a liability related to translation. First, Ssd1 is required to manage the stress of chromosome duplication, across strain backgrounds and amplified chromosomes^44^. Ssd1 has been implicated in translational repression and mRNA localization^44,72–74^, among other processes. Intriguingly*, SSD1* deletion sensitizes euploid strains to mutation of the elongator complex as well as Deg1 tRNA:pseudouridine synthase, both of which modify tRNAs to promote translational fidelity^75,76^. These links connect Ssd1 to aneuploidy and translation, but also to snoRNAs and tRNAs that are implicated in our modeling. Recent work from our lab^77^ shows that over-expression of genes involved in translation, including eIF5A protein Hyp2, complements *ssd1Δ* aneuploid growth defects. Both *SSD1+* and especially *ssd1Δ* aneuploids are inherently more sensitive to translation elongation inhibitors^24,44^, suggesting that translational stress is likely at play in wild-type aneuploids. We proposed that *SSD1+* strains can largely buffer the cost of most chromosome duplication unless otherwise compromised by translational stress^44^.

### Gene length is the strongest predictor of deleterious gene duplication

The cost of chromosome duplication is well modeled by the additive cost of duplicating individual genes on each chromosome; thus, considering the features of deleterious gene duplications can further our understanding of aneuploidy. We expected that genes encoding multi-subunit complexes and with multiple protein-protein interactions would be among the most deleterious, thus validating long-standing models of protein imbalance as a major cause of aneuploidy toxicity. However, deleterious gene duplications were not enriched for either. This recapitulates several other studies that also saw no enrichment for components of protein complexes amongst duplication-sensitive genes^53,57,78^. The absence of these signatures indicates that the Balance Hypothesis^40,79^, often invoked to explain aneuploidy toxicity, may well be true for high-level protein imbalance but not for mere duplication of genes and their native regulatory sequences. The reason is likely due to dosage control, which has been observed repeatedly for multi-subunit proteins amplified in yeast and human cells^22,53,80–84^. While some dosage control can happen at the transcriptional level^7^, much occurs post-translationally. For example, proteins encoded by human chromosome 21 show increased turnover rates^83^. Genes encoded by other aneuploid chromosomes in human cell lines also show increased degradation rates according to their role in the complex^85^. Hence, cells likely have evolved mechanisms to manage stoichiometric balance of important proteins, at least when their genes are merely duplicated^85^. One interesting observation from machine learning is that genes encoding proteins prone to aggregation (A3D prediction^86^) are less detrimental when overexpressed on the 2-micron plasmid (Fig. 6C), consistent with the idea that protein aggregation can be protective^87,88^.

We were surprised that modeling predicted a single major feature – gene length – as the strongest predictor of deleterious gene duplicates, with longer genes associated with dosage sensitivity. Remarkably, we observed that gene length was a strong predictor of dosage-sensitive genes in several screens^54,55,57,59,89^. There are several possible reasons why longer genes tend to be more deleterious when duplicated. First, gene length is correlated with multiple other biophysical features: larger proteins are more likely to contain an intrinsic disorder region, have more phosphorylated sites, and have a higher fraction of buried residues. One possibility is that gene length is simply a proxy for a multitude of other gene properties that are each mildly deleterious. However, we did not find strong support for this hypothesis: when gene length was omitted from the model, several features correlated with gene length were selected, but the model did not perform as well as using gene length alone. It remains possible, however, that longer protein primary sequences are more likely to capture some deleterious features.

Another possibility is that longer genes and transcripts create more chances for error during protein synthesis. Longer genes typically display slower translation initiation and elongation rates, a relationship conserved across organisms^90–94^. This relationship could reflect higher-order RNA structure or other features of long mRNAs^95,96^; indeed, of the subset measured, deleterious gene duplicates do have more structure (p-value = 0.0008)^97^. Longer transcripts also increase the probability of translation errors including tRNA / amino acid misincorporation, ribosome frameshifting, premature termination, and co-translational protein folding errors, all of which are influenced by sequence but also proportional to transcript length^96,98–100^. These errors in turn can lead to proteostasis stress and an energy burden to manage that stress^90,96^. Indeed, managing proteostasis stress through quality control pathways such as the Ubiquitin Proteasome System is important in sensitized aneuploid strains^101,102^; however, the direct source of the proteostasis stress remains unclear – our results suggest that translational errors could contribute.

In all, our study presents a quantitative assessment of aneuploidy cost, in a single strain and controlled environment. Although the principles reported here are likely conserved, the details including precise fitness costs of specific genes and non-genic features, as well as the generalized sensitivity of strains to translational and proteotoxic stress, could vary significantly across strains and conditions^59,103^. An important consideration for future work will be to quantify that variation.

## Material and methods

### Strains and plasmid

Strains and plasmids used are listed in Resource Table S3. YPS1009 aneuploids were generated using the methods of Hill and Bloom^45^ except Chr12 aneuploidy described in Hose et al.^44^. Briefly, a DNA cassette including the *GAL1-10* promoter (GAL1 oriented toward the centromere), HphMX6 gene for hygromycin resistance, and terminator *P_TDH3_-GFP-T_CYC1_* (except for Chr3, 9, and 16 where GFP was omitted) was integrated at 60 bp from each centromere of interest and selected on hygromycin medium. Each resulting euploid strain was grown for 16 h in YP (1% yeast extract and 2% peptone) medium with 2% raffinose and switched to YP with 2% galactose for one doubling based on optical density, and then plated for single colonies. For transformants carrying the GFP cassette, colonies were initially screened for 1X (euploid) versus 2X (aneuploid) GFP fluorescence on a flow cytometer, and colonies with 2X fluorescence were selected. Aneuploid colonies were selected via qPCR of genes on and off the amplified chromosome to confirm duplication of the amplified chromosome; selected colonies used in this study were confirmed by low-coverage whole genome sequencing, confirming that genes spanning the entire chromosome were present on average 2X higher copy than genes on all other chromosomes. *ssd1Δ* aneuploids were obtained by crossing aneuploids selected above to the euploid *ssd1Δ* and selecting resulting *ssd1Δ* aneuploid clones. YPS1009 with a duplication of Chr6 could not be generated in YPS1009, and duplication of Chr16 in *ssd1Δ* produced very sick colonies that could not be cultivated. Genomic DNA was isolated with the DNeasy Blood and Tissue Kit modified for yeast (Qiagen) and sequenced using the NEBNext Ultra II DNA Library Prep Kit on the Illumina MiSeq. Eight of the aneuploids (Chr1, 4, 5, 7, 10, 13, 14, 15 and 16) were backcrossed to remove the centromere-proximal cassette. Euploids and aneuploids with the cassette had no difference in growth rate compared to an isogenic strain without the cassette, confirming that the cassette does not influence fitness.

The pJR1 plasmid expressing 7 C/D box snoRNA encoded on Chr13 (snr72, snr73, snr74, snr75, snr76, snr77, snr78) was obtained by amplifying 2017 bp containing the polycistronic C/D snoRNAs region from Chr13 (coordinates 280,245-282,261 from the YPS1009 genome assembly) and ligating it into pJH1 plasmid. The pJR2 plasmid containing 7 H/ACA snoRNAs was obtained by ligating a fragment containing *SNR36, SNR8, SNR31, SNR5, SNR81, SNR9* (synthesized by Twist Bioscience) and *SNR35* (amplified from YPS1009) into pJH1. A fragment from the Yce1313 plasmid (shared by the Cai Lab) containing all Chr12 tRNAs was cloned into pJH1 to obtain the pJR3 plasmid. All plasmids were verified by Sanger sequencing. MAF1 was deleted by homologous recombination of the *HphMX6* cassette and verified by diagnostic PCR; aneuploid strains were generated by crossing the euploid *maf1Δ* to aneuploids.

### Growth conditions

Strain passaging was minimized to ensure maintenance of the aneuploidies. Freshly streaked colonies were used to inoculate liquid YPD and cultured for ∼1 generation before changes in optical density (OD_600_) were scored for ∼140 minutes and fit with an exponential curve to calculate growth rates. The maintenance of aneuploidy was periodically checked through diagnostic qPCR of one or two genes on the amplified chromosome normalized to a single-copy gene elsewhere in the genome (*ERV25 or ACT1*), taking ∼2X higher copy of the amplified genes to confirm aneuploidy. Detectable loss of the extra chromosome at the culture level was rarely observed, but cultures for which > 20% of final colonies reverted to euploidy were excluded from analysis. Significant differences in observed versus expected growth rate were assessed with replicate-paired T-tests. Unless otherwise noted, all studies used 4 biological replicates.

For strains transformed with plasmids (pJH1, pJR1, pJR2, pJR3), cells were cultured for 2 hours in YPD + 100ug/ml nourseothricin media then shifted to YPD without antibiotics and grown for another hour before OD_600_ measurements were collected for growth rates. The biological replicates represent the growth of at least two different transformants, transformed on different days.

### YPS1009 genome sequencing

A highly contiguous assembly of YPS1009 strain AGY731 was prepared through a hybrid approach of Oxford Nanopore (ONT, Oxford, UK) and Illumina (San Diego, California) sequencing. High molecular weight DNA was prepared for ONT sequencing by harvesting cells from an overnight YPD culture, spheroplasting, and gently lysing cells followed by phenol:chloroform extraction and ethanol precipitation of DNA. The preparation was enriched for high molecular weight DNA >1.5 kb by bead cleanup using a custom buffer (10mM Tris-HCl, 1mM EDTA pH 8.0, 1.6M NaCl, 11% PEG8000). DNA was prepared for sequencing using sequencing kit LSK-110 (ONT) and sequenced on a single flongle flow cell (ONT). ONT sequencing produced 175 Mb resulting in ∼14x coverage of the yeast reference genome. Initial base calling was done using guppy v.6.2.1 (ONT) retaining reads with Q>7. The initial assembly was done using ONT reads with Canu v.1.9^104^. This assembly was polished using Illumina data pooled from 32,723,650 reads of all YPS1009 aneuploid strains (211X YPS1009 genome coverage) using pilon v.1.23 iteratively three times^105^.

The assembly resulted in 23 contigs with sizes ranging from 1,061 to 1,482,091 bp of which 11,353,357 bp had homology to the S288c genome. Each of the 23 contigs was aligned to the S288c chromosome to which it had shown maximal homology using MUMmer^106^, with -c parameter set for each chromosome based on aligning the S288c chromosome sequence to the S288c reference genome, to minimize short off-target alignments. Four chromosomes (Chr7,12,13,16) were spanned by two contigs and one (Chr15) was spanned by 4 contigs. To evaluate alignment gaps on those chromosomes, we considered Illumina DNA read coverage from the aneuploid YPS1009 strain in which that chromosome was duplicated. We did not find support for the S288c sequence being present in YPS1009 at any of these gaps, strongly suggesting that the S288c sequence in those gaps is truly missing from YPS1009. Contigs for these chromosomes were joined by {N}_10_ representing those gaps. The final assembly resulted in 16 assembled chromosomes.

We assessed the quality of the assembly in several ways. First, the median percent identity for MUMmer-aligned segments was 99.25%, showing high similarity to the S288c genome as expected. Second, we considered the coverage of known universal single-copy orthologs from the OrthoDB database BUSCO^107^. BUSCO analysis identified 99.2% (2119 out of 2137) of the universal single-copy genes from the saccharomycetes_odb10 ortholog database, of which 2074 were in single copy and 45 were duplicated, indicating high coverage of expected genes. Base-level accuracy and completeness were measured with Merqury^108^. An optimal k-mer size (16) was generated using best_k.sh (provided by Merqury suite) and a k-mer database created with Meryl^108^. This k-mer database was used to evaluate the assembly, which returned a completeness score of 99.502%.

Finally, we annotated the gene content using Liftoff^109^. Multiple genes and other genomic elements with high level of homology were annotated to the same region, we filtered out the annotation with the lowest homology. Liftoff identified 6,552 genes, 277 tRNAs, 77 snoRNAs, 21 ncRNA, and 354 ARS in the YPS1009 genome. For transposable elements (TE), we combined Liftoff identification with ReasonaTE^110^, and collapsed TEs that were mapped to the same region. There are 23 retrotransposons containing functional GAG-POL open reading frames. Among the 6552 genes annotated by Liftoff, 57 are missing a start codon, 331 are missing a stop codon, and 70 have an in-frame stop codon.

89 genes from S288C were missing in YPS1009: 48 of them were mapped to other ORFs and filtered out, which likely correspond to genes present in multiple copies in S288C. The remaining 41 missed genes were used as BLAST queries to the YPS1009 assembly: 4 small genes aligned to multiple loci (> 8) in the YPS1009 assembly while 38 genes were not identified by Liftoff or BLAST of the YPS1009 contigs; the position of 19 of these genes in the S288C genome reside in 3 suspected gaps between YPS1009 contigs that were supported by the absence of Illumina reads mapping to those YPS1009 regions as described above. 12 genes mapped to a gap on Chr12 that was corroborated by an absence of Illumina reads. Thus, the draft assembly of the YPS1009 genome is close to complete, barring small-scale errors whose correction is beyond the scope of this study.

### Gene duplication fitness cost measurements using MoBy 1.0 plasmid library

The euploid YPS1009 strain (AGY1611) was transformed with a pool of the molecular barcoded yeast ORF library (MoBY 1.0) containing 5,037 barcoded CEN plasmids^48^. At least 25,000 transformants were scraped from agar plates for roughly fivefold replication of the library, and frozen glycerol stocks were made. Multiple independent transformations of the pooled library were performed for each strain (see competitive growth details below). Competitive growth was done in liquid synthetic media lacking histidine (SC-His) and with 100 mg/L nourseothricin and 200 mg/L G418 to maintain the plasmids. Competition experiments were performed as previously described^48,59,111,112^. Briefly, 1 mL frozen glycerol stocks of library-transformed cells were thawed into 100 ml of liquid medium at a starting OD_600_ of 0.05, then grown in shake flasks at 30°C with shaking. The remaining cells from the frozen stocks were pelleted by centrifugation and represented the starting pool (generation 0) for each strain. After five generations, each pooled culture was diluted to an OD_600_ of 0.05 in fresh media, to maintain cells in log phase. Cells were harvested and stored at −80°C after 10 generations. 7 biological replicates from 5 independent library transformations were collected and analyzed. Plasmids were recovered from each pool using Zymoprep Yeast Plasmid Miniprep II (Zymo Research D2004-A) with the following modifications: samples were incubated with 15 units zymolyase at 37°C for 1 hour, with inversion every 15 minutes; incubation in cell lysis buffer was extended to 10 minutes; after neutralization, samples were put on ice for 30 minutes, then centrifuged at 4°C. Plasmid barcodes were amplified using primers containing Illumina multiplex adaptors as described in^112^. The number of PCR cycles was reduced to 20. Barcode amplicons were pooled and purified using AxyPrep Mag beads (1.8X volume beads per sample volume) according to the manufacturer’s instructions. Pooled amplicons were sequenced on one lane of an Illumina HiSeq 4000 to generate single-end 50 bp reads. The data analysis was performed as follows: the bottom 5% of barcodes based on read abundance were removed from the total counts at generation 0, as well as barcodes that had a count of 0 at generation 0 in any sample. A pseudo-count of 1 was added to each gene in every sample in the dataset. Barcode counts were normalized using the TMM method^113^ and analyzed in EdgeR^114^ version 3.36.0 using a gene-wise negative binomial generalized linear model with quasi-likelihood tests. Results were similar when normalized by total reads per sample. Significant differences between experiment endpoint and generation 0 were defined as those with FDR < 0.05 using the Benjamini-Hochberg procedure for multiple test correction^115^. Fitness scores of 4,462 genes were calculated as the log_2_ of the ratio of normalized reads after 10 generations divided by reads at generation 0 (Table S1). Significant fitness scores are highly correlated with those from comparable YPS1009 Moby 1.0 library grown in YPD medium (R^2^ = 0.8), but not with YPS1009 transformed with the Moby 2.0 library grown under similar conditions as used here^59^, confirming that media differences between this study and Robinson *et al.* do not explain modeling differences.

### Modeling Aneuploidy fitness costs

Model 1 fits the measured growth rates (4 per strain) for each aneuploid relative to euploid cells as a function of the sum number of verified and uncharacterized genes per chromosome, according to the YPS1009 genome annotation. A total of 4,369 measured genes are mapped to the YPS1009 genome and included for further analyses. We did not consider dubious genes. Linear regression was performed using the ordinary least square (OLS) method (Statsmodels, version 0.13.5).

Model 2 fit measured growth rates described above as a function of the measured fitness costs for genes duplicated on each chromosome as follows. For measured genes that were statistically significant (FDR <0.05), the fitness cost was taken as the fitness scores described above. Genes with missing values (848 genes) or that were not statistically different from neutral (FDR > 0.05) were scored with the mean log_2_ fitness score across all measured genes = -0.33. For 624 genes that are in the collection but were not detected in our experiment, we assumed their fitness cost was too toxic to make it to the starting pool in this strain background and thus imputed values with the 2.5% lower quantile value of all genes = -3.2. Each chromosome cost was estimated based on the sum of these log_2_ values for genes on that chromosome, and the linear fit was calculated as described for Model 1. The improvement of Model 2 compared to Model 1 was estimated in two ways. First, we used a nested model and Chi-square test, considering the contribution of Model 1 (gene number) plus the contribution of Model 2 costs normalized to each chromosome’s gene number, then fitted in an OLS model. We then perform a likelihood-ratio test (Chi-Square test, degree of freedom = 1) to show that both features are significant (number of genes/Chromosome p-value: 1.2×10^-13^, normalized Chr. cost p-value: 0.045). Second, we performed 10,000 random permutations of gene fitness cost labels across chromosomes, while preserving the number of genes per chromosome in each trial and summed the permuted Chr. costs. We then fitted the aneuploid relative growth rate against every permuted Chr. cost iteration and compared the R^2^ values to Model 2 R^2^. Out of 10,000 permutations, only 4 met the observed Model2 fit for wild-type aneuploids (p = 0.0004) and none for the *ssd1Δ* strains. The importance of beneficial genes was estimated by summing detrimental/neutral genes and beneficial genes separately and fitting a multifactorial linear regression. A Chi-square test showed that both features are significantly contributing to the fit.

Model 3 was assessed by first compiling a list of non-genic features from the YPS1009 Liftoff feature detection (Table S4) and normalized to the total number of features per chromosome to prevent high correlations in between features. Features were selected using a bootstrap-Lasso approach^49^: 10000 random subsets of 60 relative growth measurements were fitted using Lasso (alpha = 0.7), and features that had a non-zero coefficient for 90% or more iteration were incorporated into a multi-linear regression model (OLS) to get model performance.

### Deleterious gene duplications classifier training

Gene biophysical features considered in the modeling are described in Table S2^44,58,86,116–131^ and available together with the gene duplication fitness costs in Table S1. Functional enrichments using GSAEpy python library^130^ (version 1.0.6) and the ontologies from Yeast modEnriChr^129^ were performed in 2 ways. First, we performed a hypergeometric test to compare genes whose duplication was deleterious (FDR < 0.05) versus the background genes set (all barcoded genes with a measured logFC). Second, we used a GSEA rank test: genes were ranked on their log_2_ fitness scores * log_10_(FDR) values. Enrichments with an adjusted p-value < 0.05 were included as categorical features for the modeling and are available in Table S1. For numerical features, a Wilcoxon rank test was performed with Benjamini-Hochberg correction^115^. To train the gene classifier to predict deleterious genes, we reduced the number of features to only those that were significant (adjusted p-value < 0.05) and removed features that were highly correlated (Spearman correlation > 0.70, see Fig. S5A), keeping the feature most strongly distinguishing detrimental genes (Fig. S5A). All models were trained and tested using a stratified 5-fold cross-validation approach: for 5 iterations, the dataset was randomly split into training and test sets while maintaining the proportion of deleterious and neutral genes. We then computed the mean and standard deviation receiver-operator curves and area under the curve (AUC-ROC) for analysis of the test set. Confusion matrices also were computed from the aggregated test set predictions. We used a seed of 17 for the k-fold splitting and all models. The following model and parameters from Sklearn (version 1.3.0) were used: Logistic regression with l2 penalty (maximum iteration = 500, solver = newton-cholesky, and balanced class weight), Random Forest classifier (n estimators = 100, minimum sample per leaf = 24, max depth = 8, minimum impurity decrease = 0.01), XGBoost Classifier (number of estimators = 100, minimum child weight = 250, subsample = 0.8, maximum depth = 4, balanced weight (0.7)), Gradient Boosting Classifier (number of estimators = 100, subsample = 0.8, minimum impurity decrease = 4, maximum depth = 6). Parameters were manually selected to reduce overfitting; Overfitting was assessed by comparing the ROC-AUC for the training and testing sets.

Models were first trained on the whole gene fitness screen from which genes with more than 6 missing biophysical features were removed (1,177 detrimental genes and 3,028 neutral/beneficial genes remaining). Due to poor predictions on the whole dataset, we focused on training binary classifiers to distinguish between medium-highly detrimental genes (log_2_ fitness score < -1.54 (quantile = 0.15) and FDR < 0.05 = 613 genes (29%)) and neutral genes (log_2_ fitness score > 0.27 (quantile 0.65), 1,472 genes). The logistic regression classifier performed better than tree classifiers or Neural networks. Features were sorted by their mean coefficients (Fig. S5B) and we observed that the 12 top features were sufficient to maintain maximal model performance with an AUC-ROC of 0.713. Features importance was assessed using a permutation feature importance strategy (Sklearn version 1.3.0, permutation_importance)^132^: each feature is randomly shuffled and the resulting degradation of the model’s score is used to compare features. Values were shuffled 10 times for each 5-fold validation dataset splitting. Feature coefficients were analyzed to assess if a feature was associated with detrimental genes or with the neutral group.

A similar classifier (Logistic regression with l2 penalty, maximum iteration = 500, solver = newton-cholesky, balanced class weight) was trained on Robinson et al. data to discriminate the commonly deleterious gene overexpression (400 genes, detrimental at FDR < 0.05 in at least 10 yeast isolates) from commonly neutral or beneficial gene overexpression (1,657 genes, not detrimental (FDR > 0.05) in at least 12 yeast isolates) measured in our lab under slightly different growth conditions^59^. In that case, no features were filtered out based on correlation.

## Data availability

The authors declare that all data supporting the findings of this study are available within the paper and its supplementary information files. The YPS1009-derivative strain genome assembly used to assign genes to chromosomes and detect non-coding genes is available on NCBI, BioProject, accession number PRJNA984736. The gene duplication screen raw sequencing and barcode counts are available on GEO (accession number GSE263221).

## Code availability

The code used to generate all findings and figures is available at: https://github.com/GLBRC/Rojas2024_Aneuploidy

## Acknowledgments

Thanks to Dr. Patrick Cai for sharing his tRNA plasmids, Michael Newton for statistical advice, and the Gasch Lab for useful discussions. This study was funded by the NIH (R01GM147271). H.A.D research is funded by an NHGRI training grant to the Genomic Sciences Training Program (T32HG002760) and an NIH training grant (T32GM007133). Research in the Hittinger Lab is funded by the National Science Foundation (DEB-2110403), USDA National Institute of Food and Agriculture (Hatch Project 7005101), in part by the DOE Great Lakes Bioenergy Research Center (DOE BER Office of Science DE–SC0018409, and an H. I. Romnes Faculty Fellowship (Office of the Vice Chancellor for Research and Graduate Education with funding from the Wisconsin Alumni Research Foundation).

## Author contributions

J.R, A.P.G: Conceptualization, Manuscript writing. J.R, J.H, H.A.D: Investigation, Methodology, Formal analysis. M.P and J.F.W: YPS1009 genome assembly, M.P. YP1009 genome annotation. C.T.H: Mentorship of J.F.W., resource sharing. A.P.G: Supervision, Funding acquisition, Project administration.

## Supplemental figures

**Figure S1.**
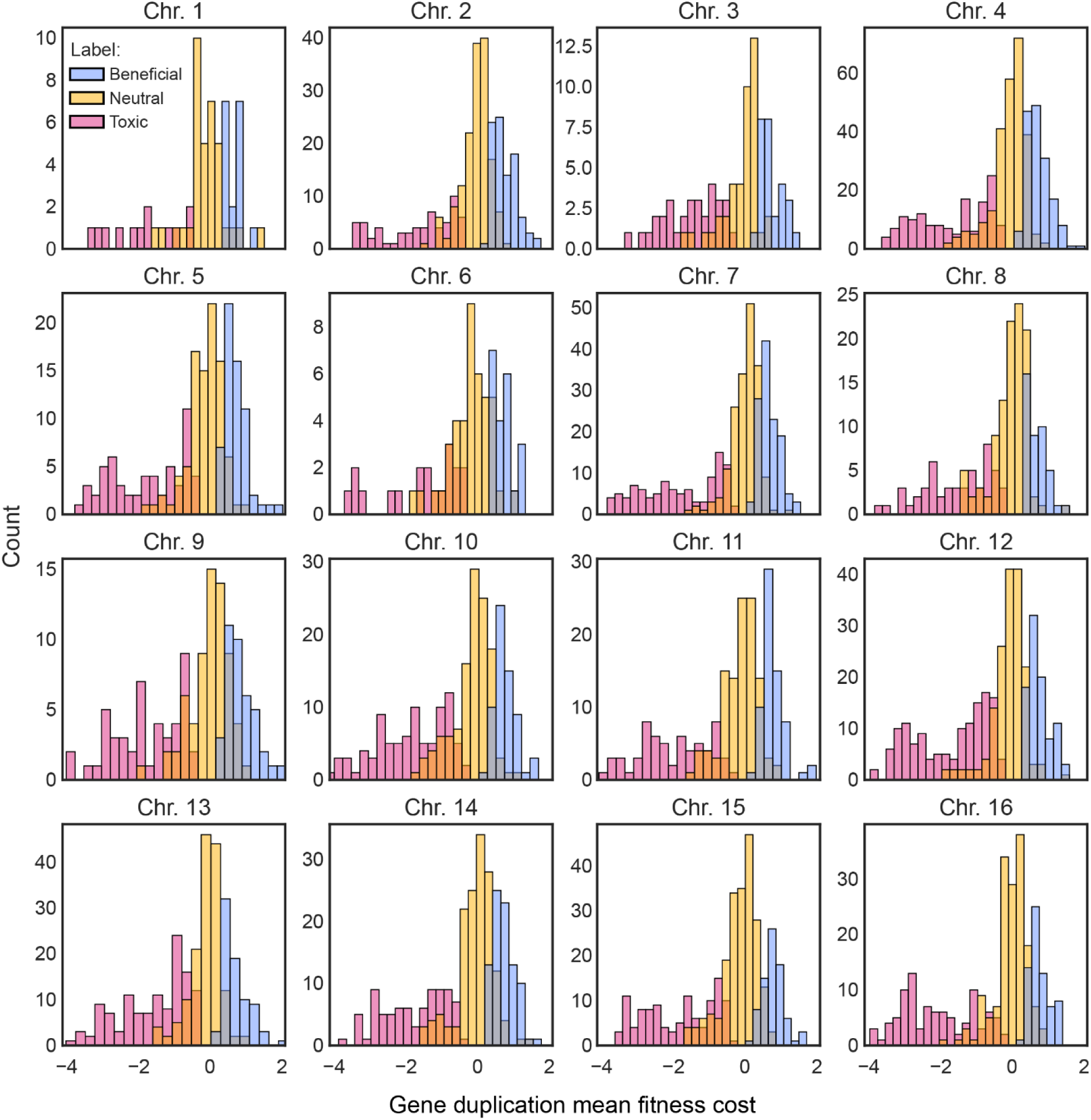
Distribution of gene fitness costs across each chromosome. Histogram of log2 fitness scores for single-gene duplications encoded on each chromosome, in each category, according to the key.

**Figure S2.**
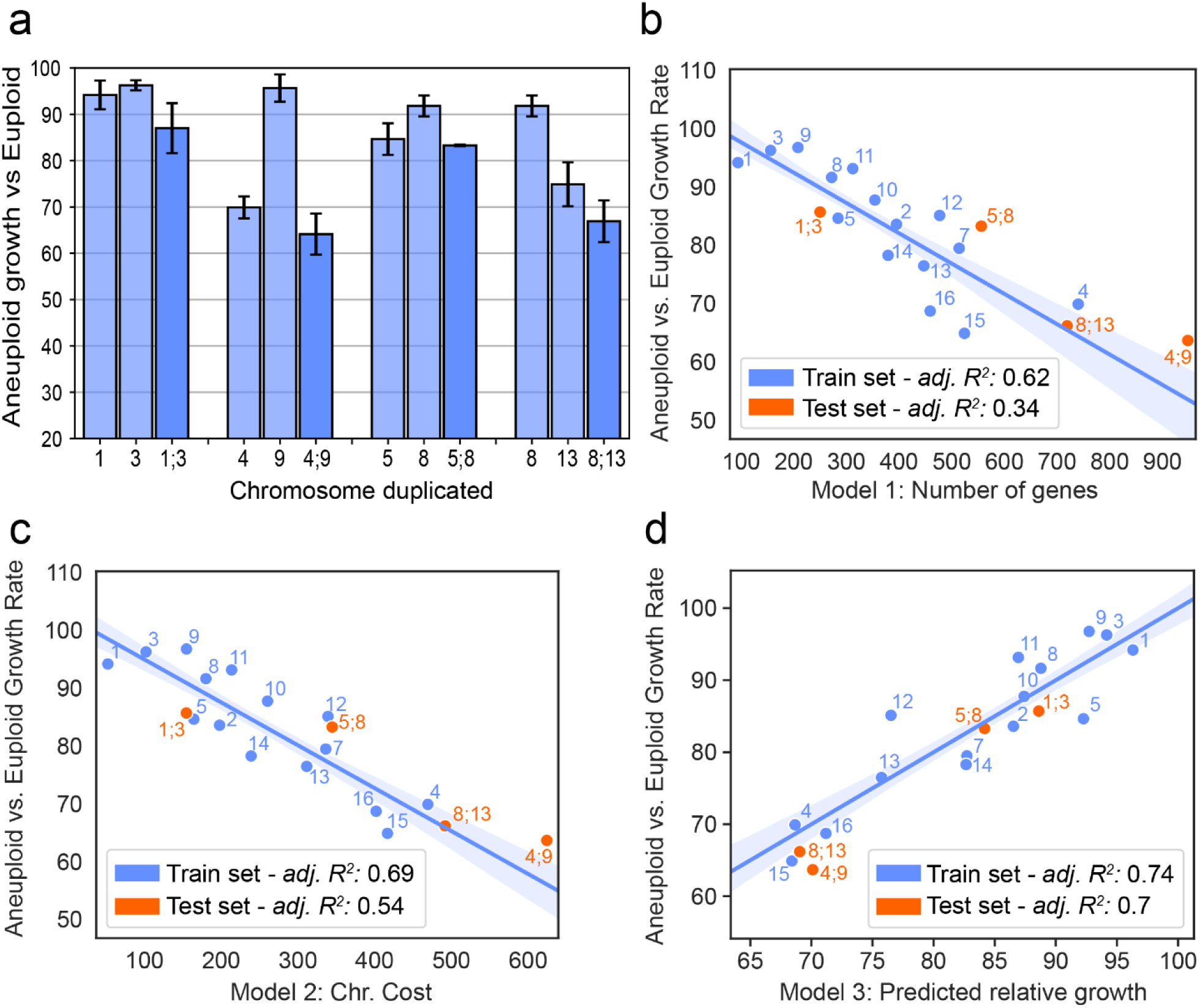
Model validation on dual-chromosome duplication strains. (A) Relative growth rates of strains with single- and dual-chromosome amplifications. (B-D) The explanatory power for dual aneuploids (orange points) is listed using the respective model trained against single aneuploids. Chromosome duplications are indicated by their number on the plots.

**Figure S3.**
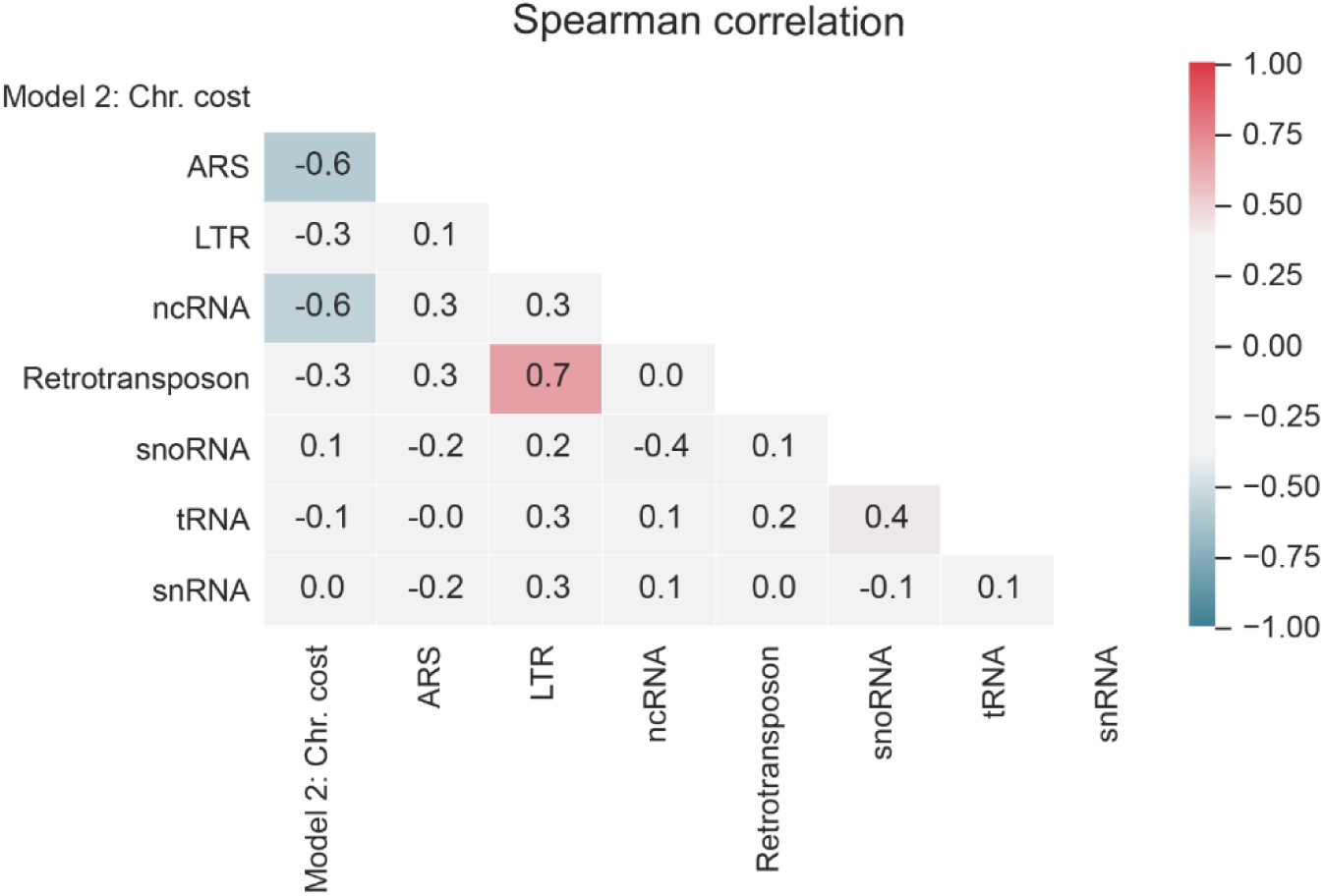
Pearson correlation analysis of chromosomal features. Except for “Model 2: Chr. cost”, the other features are normalized to the total number of features on each chromosome.

**Figure S4.**
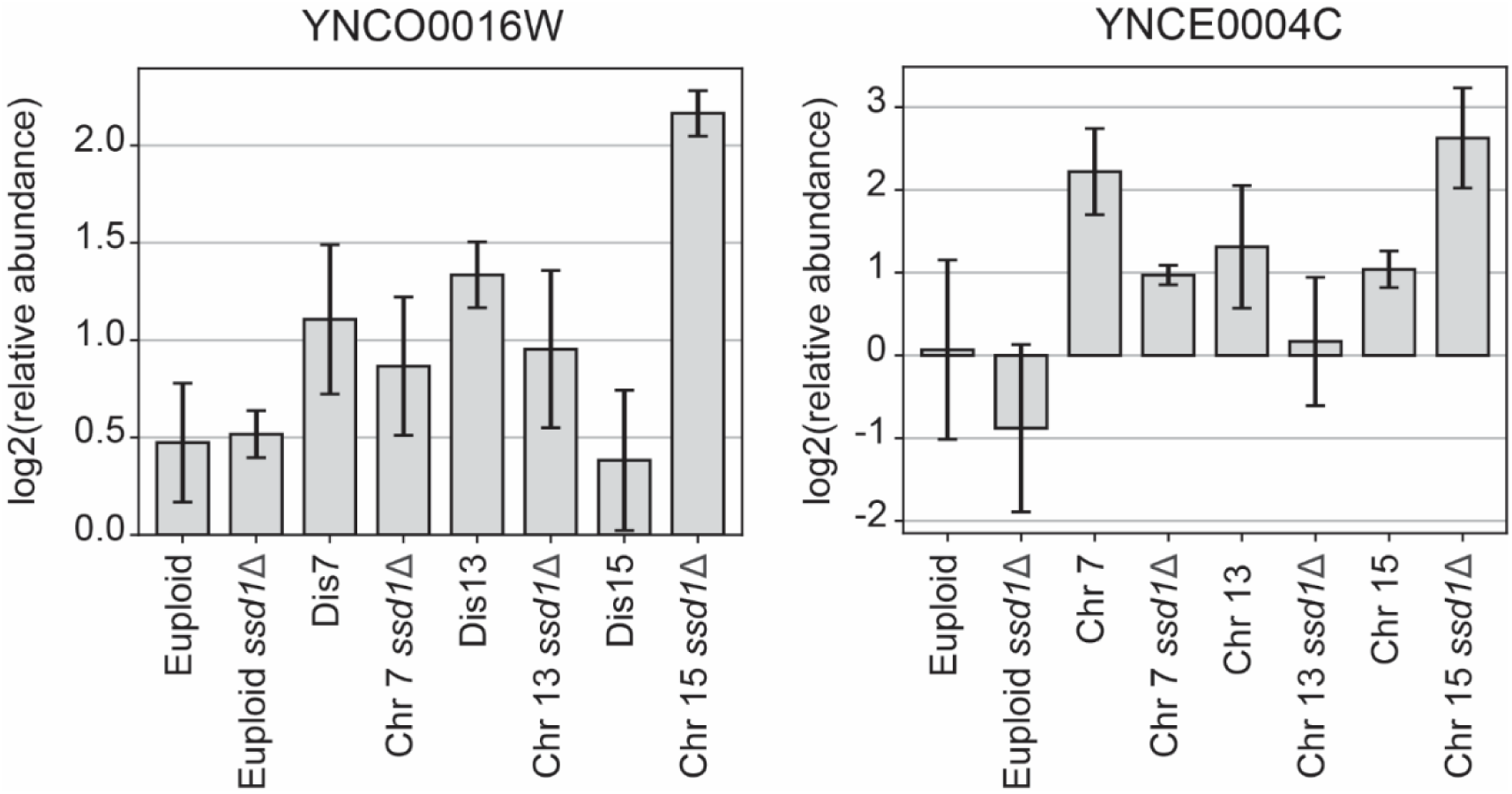
The average and standard deviation (n > 2) of the log_2_ relative abundance of two different tRNAs in *maf1Δ* versus the corresponding wild-type strains. All strains show a higher abundance of at least one (typically both) tRNA when *MAF1* is deleted. See source data for replicates values.

**Figure S5.**
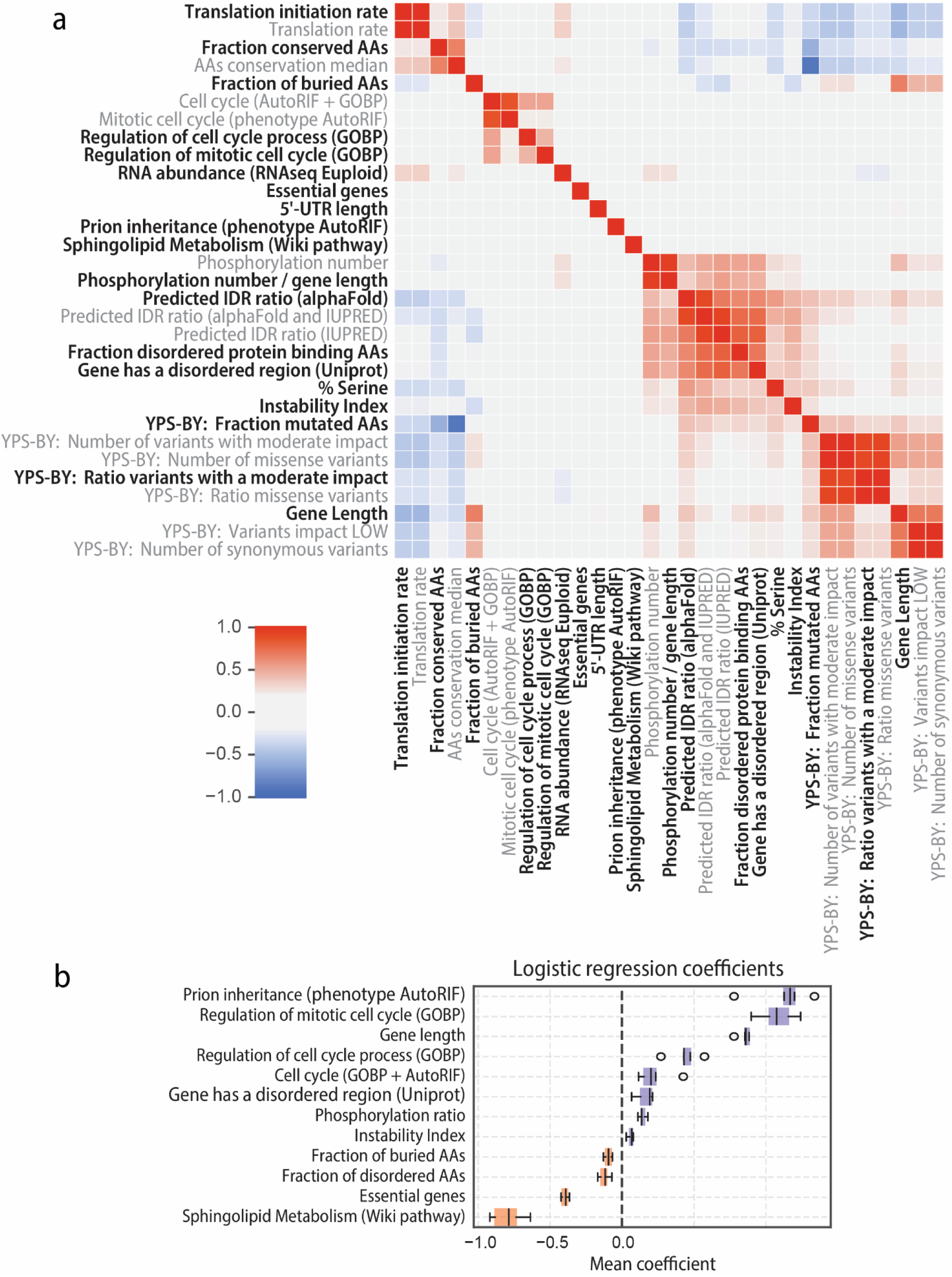
Feature selection for logistic modeling. (A) The pairwise Spearman correlation between features that significantly distinguish deleterious from neutral genes (see text), calculated across all genes in the dataset. Features correlated > 0.7 were identified and all but the most significant of those features (black text) were removed from consideration (grey text). See Table S2 for details. YPS-BY comparisons test allelic differences between the YPS1009 host strain and Moby 1.0 library. (B) Top 12 features of the logistic regression model according to absolute coefficient values (mean 5-fold cross-validation). These features were used to train the linear regression displayed in Figure 5B. In orange are features that predict neutral genes and in purple are features that associate with the group of 618 deleterious genes (filtered dataset).

**Figure S6.**
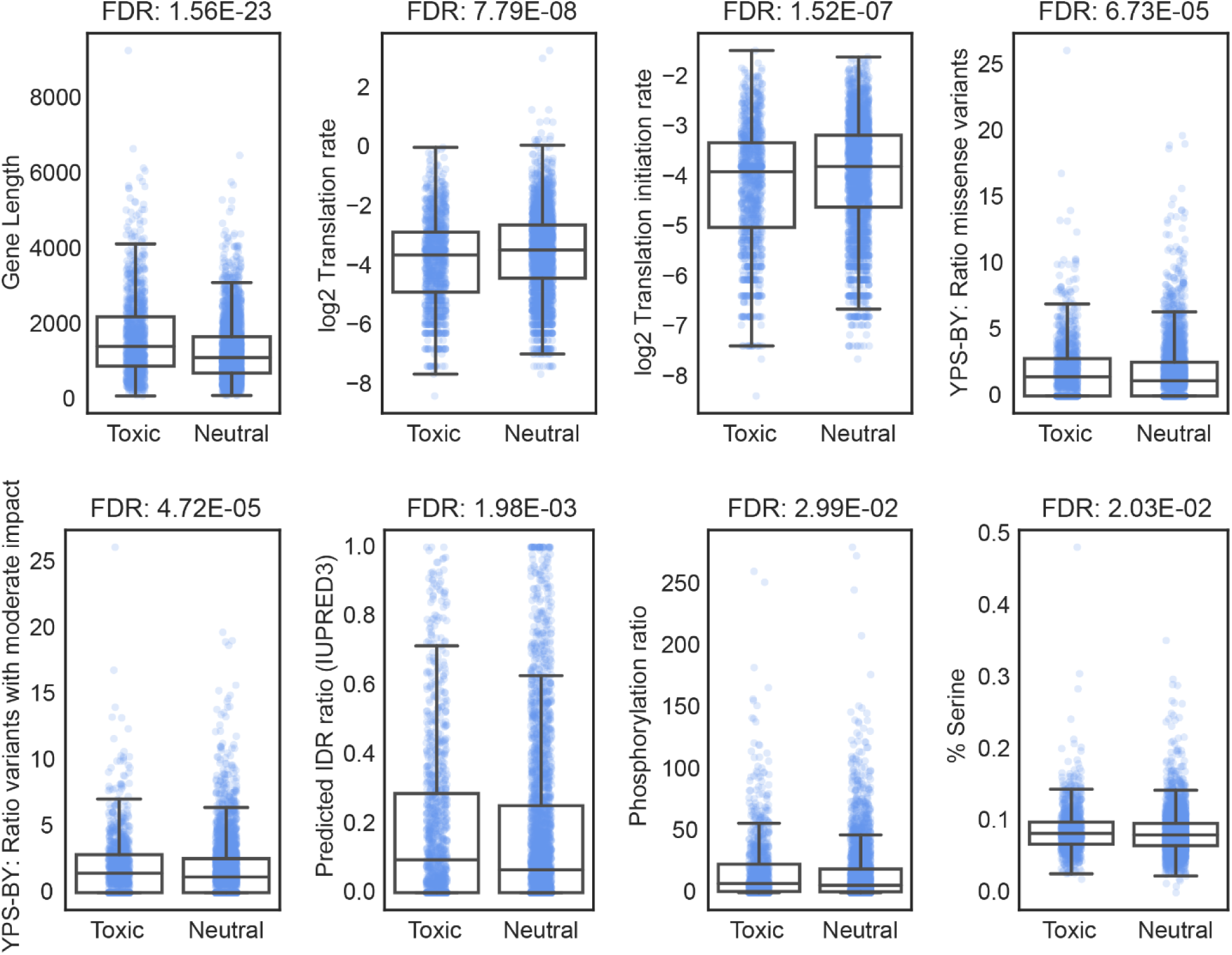
Distribution of biophysical features that distinguish duplication-sensitive (DS) genes from neutral genes. Boxplot and scatter plot of biophysical features with statistically significant differences in distribution between 1177 detrimental and 3028 neutral genes (Wilcoxon rank-sum test). For improved boxplot visualization, translation initiation and translation rate were log-transformed. FDR (Benjamini-Hochberg False Discovery Rate correction) is listed above each plot. Many of these features are correlated with gene length which is the main determinant selected by the model (see Discussion).

